# Examining food intake and eating out of home patterns among university students

**DOI:** 10.1101/320325

**Authors:** Erand Llanaj, Róza Àdàny, Carl Lachat, Marijke D’Haese

**Author notes:** These authors contributed equally to this work (Erand Llanaj, Róza Àdàny). These authors also contributed equally to this work (Carl Lachat, Marijke D’Haese).

## Abstract

Eating out of home (OH) is increasingly popular in Balkan countries, among them Albania. To date there is only anecdotal evidence regarding nutritional quality of food consumed OH and the contribution to diet. This study assessed intake of foods and drinks consumed OH and at home (AH), as well as their nutritional contribution to the daily diet of university students in Tirana, Albania. Using a single day Automated Multiple Pass Method (AMPM) 24-hour dietary recall, we examined food intake among 289 students aged 18-24 years old, from three major universities in Albania. Contribution of eating OH to total energy intake per day, as well as to daily consumption of macronutrients by eating OH intensity tertiles were assessed. Foods and drinks consumed OH contributed 46.9% [95%CI:41.4-52.8] of total daily energy intake, representing, on average, 1169.1kcal [95%CI:1088.3-1249.9]. Sweets, soft drinks and meat products were more frequently consumed OH, while fruits and vegetables consumption was extremely low. The average quantity of sugars and dietary fats per day was higher AH, 76.9g [95%CI:70.3-83.5] and 173.7g [95%CI:163.2-184.2] respectively, compared to OH, 33.7g [95%CI:30.4-37.0] and 142.0g [95%CI:131.5-152.5] respectively. Dietary composition of AH intake was richer in sugars, total fats and proteins, while OH intake was richer in saturated fats. The overall diet appeared unhealthy, when nutrients were assessed as energy percentage against WHO proposed nutrient standards for sugar and saturated fats. Eating OH, even though was associated with lower fruits and vegetables intake, was not clearly associated with poor diet quality, as AH foods were also characterized by increased saturated fats and sugars intake as energy percentage. This study provides data on the first assessment of current dietary patterns of the studied population and can be used as baseline for designing and conducting future studies and interventions targeting malnutrition in all its forms.

## Introduction

Since its transition from a central planned to a market-oriented economy in early 1990s, Albania has experienced economic growth, fast urbanization and regional trade liberalization, which are some of the major drivers of ‘westernization’ of diets: increased intake of meat, fat, processed foods, sugar and salt [1]. The transition to an upper middle-income country led to an increase in supply and availability of food and prevalence of overweight and obesity has steadily increased to a recent estimate of 58% in males and 45% females [2]. There is a clear urgency for evidence on diet and food consumed in the country, which is currently absent, as noted from a recent review of nutritional survey in 53 European countries of the WHO European region, including Albania [3]. Such evidence is yet needed to aid development of effective policy measures and interventions. Food service providers, such as cafes and fast-food vendors are popular in the Balkan countries and are increasingly important sources of foods and drinks for Albanian consumers [4]. Between 1994 and 2000, a tenfold increase was recorded for expenditures on eating OH, attributed to urbanization and high availability of relatively inexpensive fast-food shops [5]. It is typical to find kiosks and fast-food vendors around schools, among them universities, generally selling energy-dense foods, while at the same time regulatory and informative measures for the nutritional quality and safety of these foods is generally poor [6, 7]. The available fast-food items are mainly meat-based wraps, pizzas, chips, soft drinks, croissants, processed meat products and different types of sandwiches, characterized by high macronutrient density and degree of processing.

Moreover, consumption of fast foods, which is associated with a westernized diet, seems to impair immunity in the long term. Evidence shows that even long after switching to a healthy diet, inflammation towards innate immune stimulation is more pronounced. These long-term changes may be involved in the development of arteriosclerosis and diabetes, diseases later in life [8]. Concurrently, food environments play an important role and pose a great opportunity to address malnutrition challenges in all their forms [9, 10].

Previously, the consumption of foods prepared OH has been linked to increased energy intake [11], increased body mass index (BMI), among those that consume these foods [12] and prehypertension in young adults [13]. Eating OH has also been associated with a decreased intake of fruits and vegetables [14], high energy density of food [11] and a lower intake of micro-nutrients [15]. Moreover, evidence suggests that there is a strong association between living in areas with increased exposure to fast food and fast food consumption as part of eating OH [16].

Studies conducted in the United States and in Brazil showed that OH foods contained less quantities of protein, iron, calcium and vitamin A and more sugars and fats in comparison with foods consumed AH [17, 18]. Investigating food intake and dietary habits in early adulthood is a critical predictor of nutritional and health status in later stages of life [19, 20]. At the same time, the transition from adolescence to young adulthood and eventually to adult life, offers an important window of opportunity to prevent risk factors for diet-related noncommunicable diseases later in life [21]. To date however, the evidence on eating OH and AH contribution in diet and dietary patterns in Balkan countries, particularly in Albania remains colloquial and little or no quantitative food intake data is available. Against this background, this report aims to fill the gap in knowledge and to assess the importance of eating OH and AH practices amongst students in particular. The study documents foods and drinks, cost, macro-nutrient and energy intake and its association with eating OH and AH patterns among university students in Tirana. Nutritional studies, like this one, intend to help inform current efforts, interventions, professionals and policy makers on the potential effects of eating OH in early adulthood in Albania and other countries in the Balkans that share similar nutritional contexts and challenges. More importantly, efforts to advance and raise nutrition profile can be coupled with the current agenda of the United Nations Decade of Action on Nutrition (2016-2025) which is a global response to nutrition challenges [22].

## Materials and Methods

### Definition of eating out of home

Eating OH may be defined by either the place of consumption or source of food. In the literature, ‘eating out of home’ and ‘away from home eating’ tend to be used interchangeably. Both concepts refer to the same notion of practices, involving foods and drinks prepared OH.

In the present study, we considered OH foods to include all foods that were not prepared at home and were obtained near fast foods, restaurants, street food vendors and other OH sources of food, including food products purchased ready-to-eat from food stores, such as supermarkets, convenience stores and some special food market. Meals were determined through the time of the day foods and drinks were consumed, and based on this, subjects were asked to classify the meal in the questionnaire as breakfast, brunch, lunch, snack, dinner or late-night meal.

### Study design

Data were collected using a cross-sectional survey in the three largest universities in Albania, namely University of Tirana, University of Medicine Tirana and Polytechnic University of Tirana. In 2015 and 2016, 36.4% and 37.8%, respectively, of all bachelor students in Albania were studying in one of these universities [23]. Data were collected between October and December 2015 and between January and February 2016. The universities involved had seven, five and six faculties, respectively. Cluster sampling was used to select individuals. The following eligibility criteria were used for inclusion of subjects in the study: (i) being enrolled in one of the three universities and (ii) studying in any of the Bachelor programs. The sample size of this study was calculated to be able to detect difference in energy intake based on nutrition research sampling guide [24]. Sample size has been used in similar study design investigating changes in energy intake AH and OH [25]. In this study sample, female population was predominant and this is not a sampling or selection bias, rather it is a reflection of the actual sex proportions of student population [23].

Height and weight of the participants were measured by trained interviewers in duplicate. A third measurement was performed if first two measurements differed by ≥ 100 grams for weight and ≥1 cm for height. The average of these measurements was taken into account and used for later analysis. BMI categories were defined based on classifications performed before in similar studies [25]. The 24-hour dietary recalls were conducted by using the AMPM [26], which uses a standardized wording methodology to make recall of all possible foods as accurate as possible. Students of the Master of Public Health, at the University of Medicine Tirana were recruited and trained on the use of 24-hour AMPM instrument. This procedure enabled them to generate more reliable data and to detect possible biases related to data collection. Each member of the team was monitored in real time by the coordinator and detailed information on the food consumed was scrutinized in order, to avoid potential missing consumption due to unawareness or unasked questions.

24 hours AMPM involves five consecutive steps: (i) listing of foods consumed by the respondent in a 24-hour period on the day before the interview, (ii) additional recall of foods by focusing respondent’s attention on nine categories of foods that are often forgotten: non-alcoholic beverages, alcoholic beverages, sweets, savoury snacks, fruit, vegetables, cheeses, bread and rolls, and any other foods, (iii) assessment of time when the food was consumed and the name of the eating occasion, (iv) collect a detailed description of each food reported, and (v) final opportunity to recall foods (Fig. 1).

**Fig. 1.**
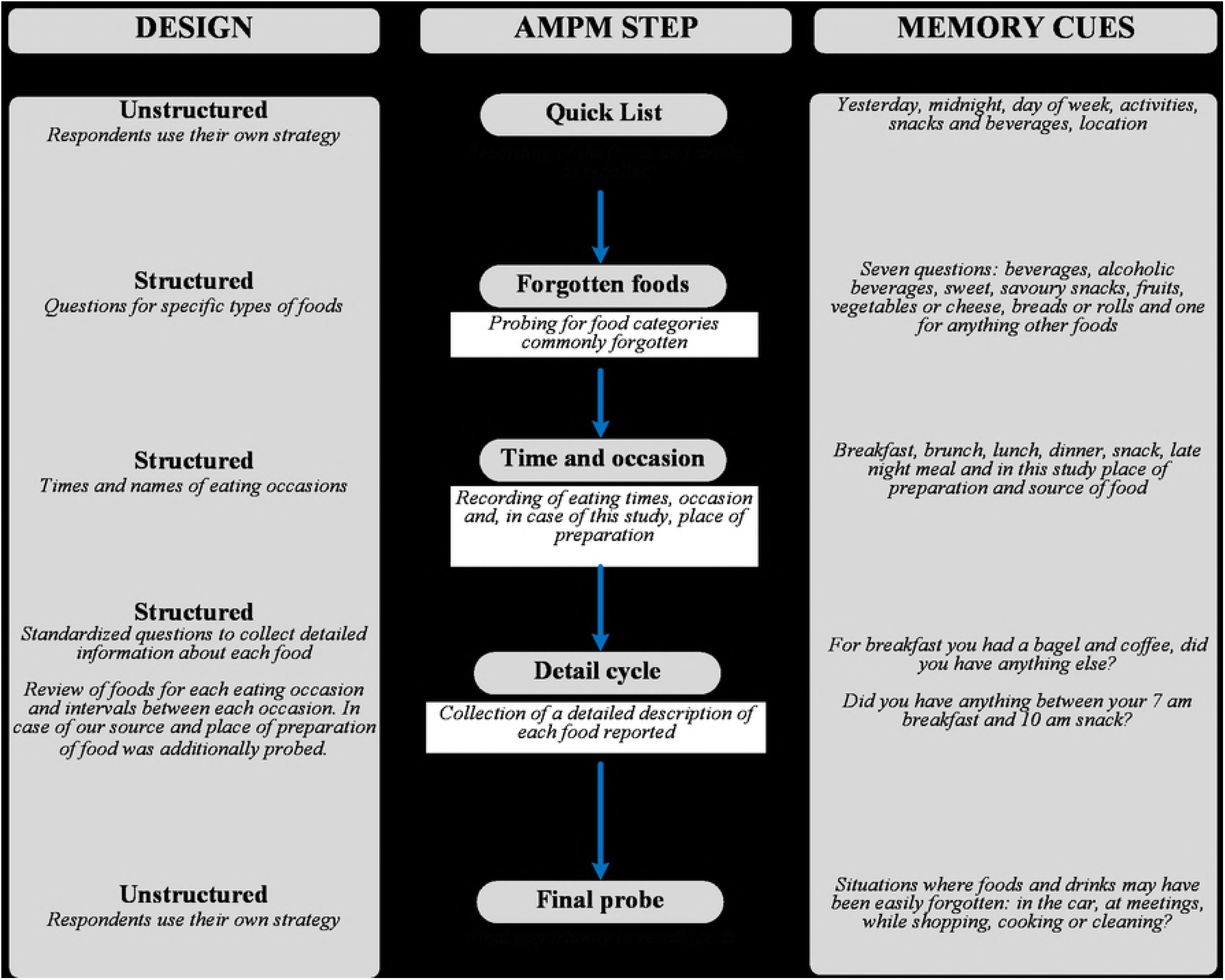
Automated Multiple Pass Method steps used in the study.

Source: Adopted from Steinfeldt et al. (2013), doi: 10.1016/j.profoo.2013.04.022

For more accuracy, a booklet with life-size drawings of different containers and dishes facilitated the whole interview process in order to make it easier for the subjects to quantify the amount of food and drinks consumed. Moreover, food recording started on different weekdays, in order to record eating patterns across different days of the week. All included subjects reported that their intake was of a typical day, in particular that: (i) intake of last 24-hours represented what they typically consumed, (ii) there was no special event (birthday, party, etc.) prior the recall, (iii) they have been consuming more or less the same foods for the last 10 days, (iv) they had no disease prior or during this intake, and (v) they were not on a diet of any kind.

The study protocol and methods were approved by UZ Ghent, Medical Ethics Committee with study number EC/2015/1118. This study was also approved by the Ministry of Health in Albania (Directorate of Health Care) and also the Ethics Committee of the University of Medicine Tirana.

Food intake data were entered and processed using the Lucille Food Intake application, similar to previous studies [27]. Outliers in dietary assessment were kept, as they did not have any impact on the results, after exploring with a sensitivity analysis. To assess the nutritional contribution of OH foods, three groups of the energy contribution of OH foods were prepared based on the tertiles of percentage energy contribution from eating OH on a daily basis. Tertiles were determined by dividing the OH energy contribution in three categories: (1) low (everything below 25^th^ percentile), (2) moderate (everything between 25^th^ and 75^th^ percentile) and (3) high (everything above 75^th^ percentile). Appropriate tests considering the sampling method were used (t-test and trend analysis) and all assumptions for normality were checked.

Statistical analyses were performed with SPSS ver. 21 (Statistical Package for Social Sciences, Chicago, Illinois), and R Statistics/R Studio. This paper has reported results in accordance with STROBE (STrengthening the Reporting of OBservational studies in Epidemiology) extension for nutrition and dietary assessment [28].

### Food composition and food cost data

Similar to other Balkan countries, a food composition database is not yet established in Albania. For the purpose of this study the Greek food composition tables were used [29]. These tables are comprehensive and perhaps, the most complete food composition tables in the region. At the same time Greek dietary patterns are very similar to the Albanian diet [30]. Data on food intake, foods and beverages items was coupled with price data of each individual food item and their weight. Prices in Albanian LEK (ALL) were obtained from local fast-foods, supermarkets, restaurants and other food vendors around the university facilities where the data collection was performed. The final result was used to compare cost of foods and drinks consumed AH and OH.

## Results

### Differences in consumption patterns

In this study, initially 364 subjects were invited to participate. A total of 35 participants reported that the recall day was not representative of a typical day and were excluded. From the 326 remaining participants, 11 were excluded due to disease (flu, common cold, etc.) during the past 24 hours. An additional 26 refused to participate later in the study and were not included in the analysis. A priori high and low sex-specific cut points for energy intake were established and 3 cases that fell outside of the cut points were excluded. Eventually 289 participants were included and considered during data analysis. Of the final sample included (87.2% females), 33.6%, 45.7% and 20.7% were students from the University of Tirana, University of Medicine Tirana and the Polytechnic University of Tirana, respectively. Participants were on average 19.7 years of age (Table 1). About 11.1% and 1.4% of the participants were overweight and obese respectively. Average daily energy intake was 2492.8 kcal [95%CI: 2427.4-2558.2].

**Table 1.** Characteristics of study subjects (N=289)

| Variable | Females |  |  | Males |  |  | Overall |  |  |
| --- | --- | --- | --- | --- | --- | --- | --- | --- | --- |
|  | Mean | 95% CI |  | Mean | 95% CI |  | Mean | 95% CI |  |
|  |  | Lower | Upper |  | Lower | Upper |  | Lower | Upper |
| Height (cm) | 164.2 | 163.4 | 165.0 | 176.7 | 176.0 | 177.3 | 165.8 | 164.7 | 166.6 |
| Weight (kg) | 56.9 | 56.0 | 57.9 | 74.0 | 73.1 | 75.0 | 59.1 | 57.9 | 60.3 |
| BMI (kg/m<sup>2</sup>) | 21.1 | 20.8 | 21.5 | 23.8 | 23.4 | 24.1 | 21.4 | 21.0 | 21.7 |
| Energy OH (kcal) | 1079.7 | 1000.0 | 1159.4 | 1752.6 | 1521.8 | 1983.4 | 1165.2 | 1059.6 | 1222.0 |

95%CI: 95% Confidence Interval for the mean

Eating OH provided 46.9% [95%CI:41.4-52.8] of the total energy intake per day. Males consumed a higher percentage of OH energy compared to females (52.9% vs. 35.9%) on a daily basis. The mean energy contribution of eating AH was higher compared to OH (Table 2). AH foods and drinks had a higher total fat and carbohydrate contribution on a daily basis, but this difference was not statistically significant for the fats consumed.

Sodium intake was similar in both categories. Most of the sugar consumed was provided by AH foods and drinks. OH foods and drinks provided on average 17.9 g [95%CI: 16.3-19.5] of saturated fats, while intake from AH foods and drinks was 15.0 g [95%CI: 13.9-16.1] on a daily basis.

**Table 2.**
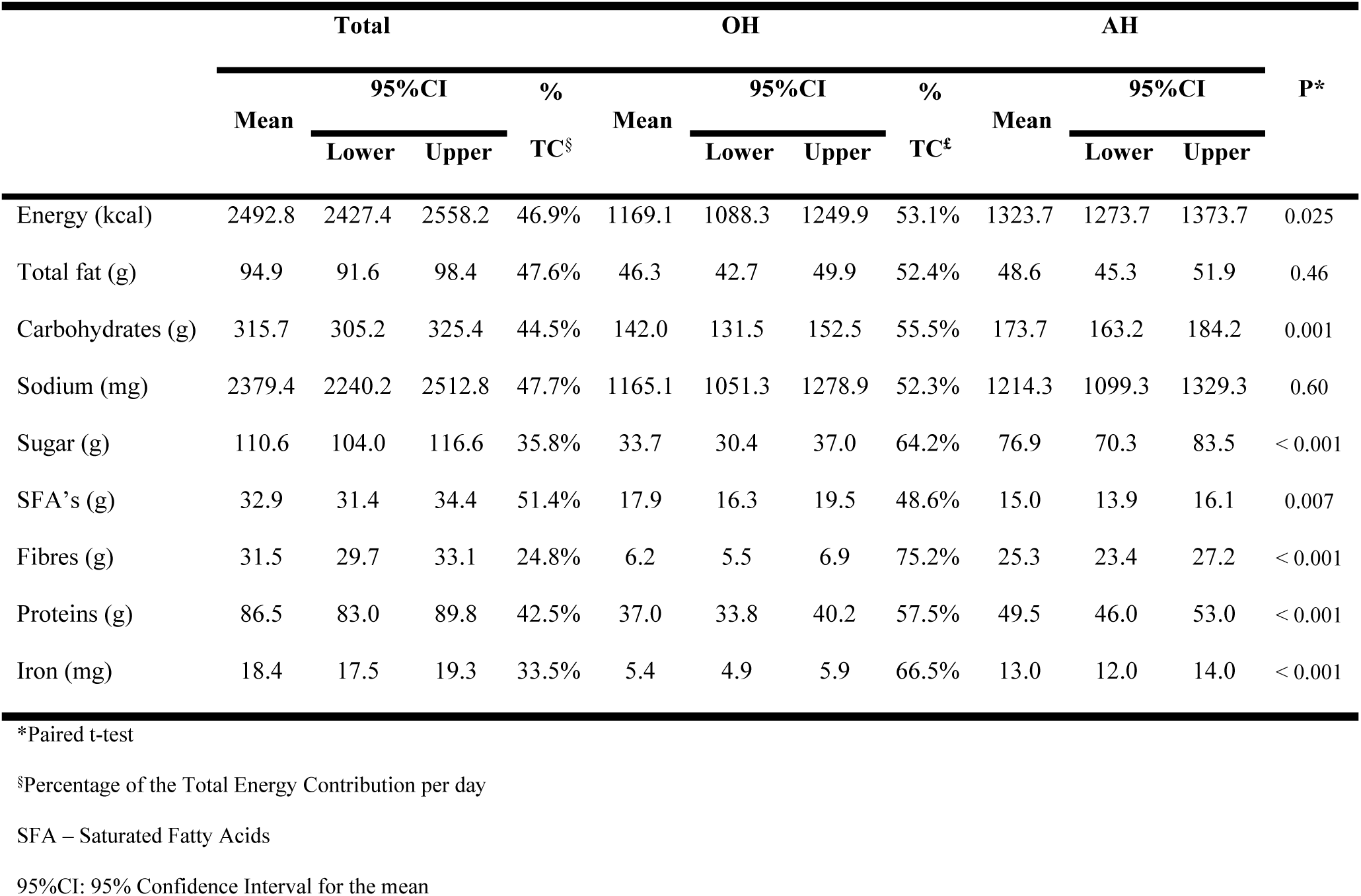
Energy of macro- and micro-nutrient intake for OH and AH foods and drinks per day.

When examining for food items ranked according to their energy intake, fruits and vegetables, meats, poultry, fish, eggs, nuts, seeds and milk and milk products were mostly consumed AH, in contrast to sweets and drinks (see S1 Table and Fig. S1).

The intake of cereal products was similar AH compared to OH, while intake of sweets and drinks was considerably higher OH (80.2% of sweets and drinks items consumed). Yoghurt was mainly consumed as homemade yoghurt, a common practice in Albanian households. Traditionally prepared yoghurt is still more consumed (81.6% of items consumed) compared to commercially prepared yoghurts (18.4%).

### Contribution of OH consumption to intake of nutrients

Table 3 shows the differences of food and macronutrient density for different intake intensities of OH foods per day. The nutrient density of fats, saturated fats, cholesterol, carbohydrates, proteins, sodium and energy per gram increase when eating out of home contribution significantly increases. On the other hand sugars and dietary fibres per gram significantly decrease. Iron (mg per gram) seems to decrease as well, however this is not a statistically significant trend. This pattern reaffirms previous observation on low consumption of fruits and vegetables and substantial consumption of fatty foods OH.

**Table 3.** Nutrient density by tertiles^§^ of energy contributed from eating OH.

|  | Low OH<sup>†</sup> | Moderate OH | High OH | P* |
| --- | --- | --- | --- | --- |
| **Energy (% OH)** | 27.0% [22.0-32.6]• | 52.0% [46.0-57.8] | 71.0% [65.2-76.0] | - |
| Energy (kcal) | 374.129 [329.625-418.633] | 1054.810 [1008.785-1100.835] | 2080.696 [1953.879-2207.512] | <0.001 |
| Energy (kcal/ gram) | 1.476 [1.406-1.546] | 1.616 [1.567-1.665] | 1.855 [1.773-1.936] | <0.001 |
| Total fats<sup>‡</sup> | 0.057 [0.052-0.063] | 0.064 [0.060-0.068] | 0.073 [0.069-0.077] | <0.001 |
| Saturated fats | 0.018 [0.016-0.019] | 0.022 [0.021-0.023] | 0.027 [0.025-0.029] | <0.001 |
| Proteins | 0.049 [0.046-0.053] | 0.060 [0.057-0.064] | 0.063 [0.058-0.067] | <0.001 |
| Carbohydrates | 0.196 [0.185-0.206] | 0.206 [0.199-0.213] | 0.238 [0.224-0.251] | <0.001 |
| Sugars | 0.086 [0.077-0.095] | 0.070 [0.065-0.075] | 0.067 [0.060-0.074] | 0.001 |
| Dietary fibres | 0.025 [0.023-0.027] | 0.020 [0.019-0.021] | 0.019 [0.017-0.020] | <0.001 |
| Sodium (mg/per gram) | 1.269 [1.117-1.421] | 1.659 [1.527-1.791] | 1.816 [1.638-1.993] | <0.001 |
| Iron (mg/gram) | 0.013 [0.012-0.014] | 0.012 [0.012-0.013] | 0.011 [0.010-0.013] | 0.107 |
| Cholesterol (mg/gram) | 0.132 [0.108-0.155] | 0.164 [0.148-0.181] | 0.170 [0.152-0.187] | 0.030 |

<sup>§</sup> Categories of eating OH contribution was calculated based on tertiles
† % of energy intake OH per day
\*Trend analysis
• Value (95% Confidence Interval for the value: lower and upper bounder)
‡ Every value is expressed in gram/per gram unless otherwise indicated
OH: out of home

When energy intake is compared from OH and AH consumption by meal (Table 4), of interest is the high-energy contribution of snacks, dinner and late-night meals consumed OH. The dinner and late-night meals contribute substantially to fat and sugar consumption compared to other meals (see also S2 Table). Differences between meals are obvious and variability of energy contribution is higher in case of meals eaten OH compared to those consumed AH.

**Table 4.** Distribution of average daily intake expressed as percentage of total and mean in kcal by meals prepared AH and OH.

|  | OH |  |  |  | AH |  |  |  | P* |
| --- | --- | --- | --- | --- | --- | --- | --- | --- | --- |
|  | % TE<sup>‡</sup> | Mean (kcal) | 95%CI |  | %TE | Mean (kcal) | 95%CI |  |  |
| Breakfast | 16.0% | 431.2 | 381.6 | 480.8 | 14.3% | 313.8 | 276.7 | 350.9 | <0.001 |
| Brunch | 16.9% | 454.2 | 402.5 | 506.0 | 8.3% | 181.4 | 133.4 | 229.5 | <0.001 |
| Lunch | 21.2% | 568.7 | 494.9 | 642.4 | 31.3% | 686.3 | 639.5 | 733.1 | <0.001 |
| Snack | 17.0% | 456.1 | 394.5 | 517.7 | 26.0% | 570.6 | 522.5 | 618.6 | <0.001 |
| Dinner | 13.8% | 370.7 | 276.7 | 464.6 | 8.3% | 182.2 | 139.0 | 225.5 | <0.001 |
| Late night meal | 15.1% | 406.3 | 343.6 | 468.9 | 11.9% | 260.9 | 220.3 | 301.6 | <0.001 |

\*Paired t-test (for mean energy density/meal)
‡ Contribution to average energy intake per day
OH: out of home; AH: at home
95%CI: 95% Confidence Interval for the mean

### Cost of eating out of home and at home

After comparing price of foods and drinks eaten AH and OH, it seems that OH food items cost on average 33.1 LEK (~0.25 EUR) more than AH ones. Further, subjects spent significantly more when they choose grains, meat products, dairy and sweets and drinks OH compared to AH. However, the cost of fruits and vegetables did not differ between AH and OH preparation (Fig. 2).

**Fig. 2.**
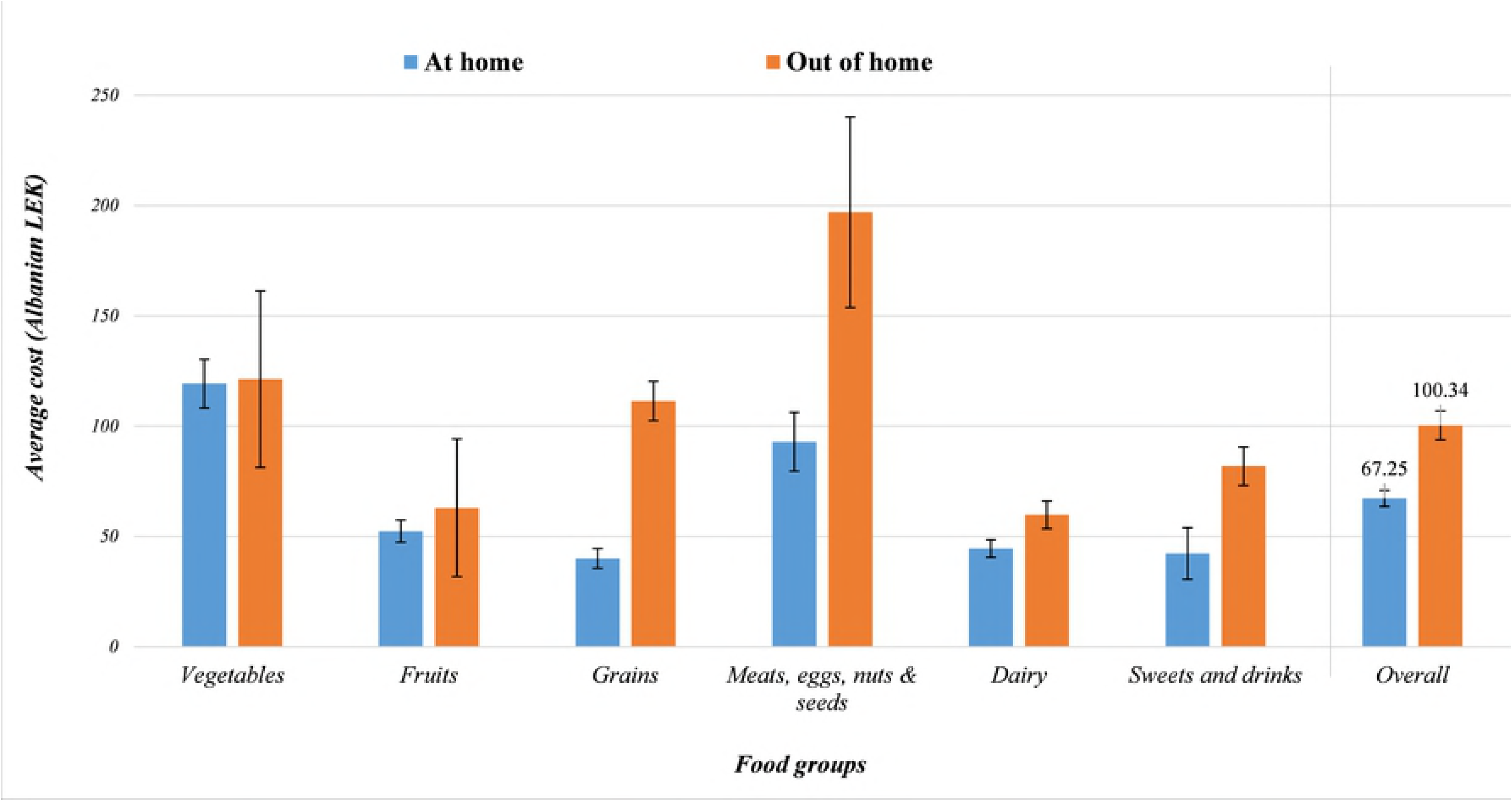
Cost of food by place of preparation and food group

## Discussion

This study sought to improve understanding of the quickly evolving eating OH patterns in Albania, which are usually associated with greater energy contribution [12, 31]. Although many studies show that eating OH is commonly associated with unfavourable dietary intake, our data indicate this is unclear in the case of Albanian students. Influencing factors or potentials of OH eating in different domains are to be investigated or reported. AH foods consumed contain higher amounts of sugar and carbohydrates compared to OH intake, perhaps due to traditional high consumption of bread, particularly white bread, spread with jam and different energy-dense spreads, containing large amounts of added sugar (e.g. chocolate spreads, honey, other sweet spreads, lard, etc.). Most OH eating occurs at fast food vendors found around universities.

No major observable differences between foods consumed AH and foods consumed OH are shown by our results, even though carbohydrate, sugar and protein amounts seem higher AH and saturated fats amounts are higher OH. Yet, important differences are noted in types of food consumed OH and energy densities of these foods. Fruits and vegetables were significantly consumed more AH, which is consistent with other studies, and were scarcely present in OH intake [12, 31] (see also Fig. S1 and S1 Table). The larger contribution of carbohydrates, sugars, fibres and proteins from AH foods, is believed to be a result of the limited food options, even AH.

Students seem to have few options, because sources of foods and food choices themselves are relatively limited. OH foods contained higher amounts of saturated fats, in terms food composition, but the contribution of AH foods for saturated fats was higher in terms of energy percentage. Although AH foods were an important source of fats consumed, compared with OH foods, the difference was statistically irrelevant, both in absolute nutrient content and contribution as energy percentage.

Sodium contribution from OH foods was high, but also for both AH and OH it was below the WHO/FAO/UNU and EU standards. Salt appeared to be consumed in relatively moderate amounts both AH and OH, without exceeding WHO/FAO proposed intake limits. This pattern, however, does not apply for other micronutrients. Although there was no clear pattern of micronutrient presence in AH and OH foods, as well as for distributions of micronutrients among AH or OH food intake, it is interesting to notice a consistent pattern regarding micronutrient presence among males and females. In all sets of micronutrients females have consistently a higher intake of micronutrients compared to males, regardless of the source of food (AH or OH). This persistent pattern of males having consistently poorer micronutrient content in recorded intake, can suggest a poorer diet in micronutrients among male subjects, especially OH, (see Figs. S3 to S5). However, due to the limitation of the single 24 hour recall and the lower representation of males in the sample we cannot infer confidently that this may the case.

The absent statistical differences between carbohydrates, fats and proteins as percentage of energy intake, consumed AH versus OH, suggest that foods consumed AH are not substantially ‘healthier’ compared to foods consumed OH in our sample. However, it is noticed that nutrient density increases for higher OH consumption (when compared with OH consumption intensity tertiles) and OH foods seem less healthy with higher energy density, sodium, saturated fats and cholesterol per gram of food consumed. Moreover, both AH and OH foods and drinks consumed have high energy percentage contribution for the 10% of daily energy intake as recommended by WHO/FAO/UNU, let alone when applying a lower threshold of 5% for additional health benefits (see Fig. S2 and S3 Table).

The ambiguous differences between AH and OH foods and drinks may be speculated to be attributed to the food environment matrix and regulatory, legal and food market infrastructure. Studies in other school settings show that availability of healthy food options influence nutrition quality and nutrition status [32, 33]. Results from this study may suggest that students have developed certain eating patterns of rather unhealthy foods and drinks, rich in sugar and unsaturated fats which is consistent AH and OH. Availability of food based on dietary guidelines in canteens are an effective way to ensure healthy diets in students [34]. However, to date, campus or university student restaurants or canteens are not available and this poses a great opportunity to consider this particular aspect of the food environment. Coupled with regulatory and voluntary policies to set standards for nutritional content and quality of meals offered at student restaurants in the future, may be a promising and beneficial with regard to public health nutrition.

Instead, students currently are limited to snack-bars offering sugar-sweetened beverages and healthy food choice has become an alternative, not a baseline. Therefore, they are pushed to eat out whatever may be available in the fast food shops and street food vendors in their reach. Fast foods vendors cluster around student residencies and neighbourhoods where students usually live and interact. Additionally, from systematic evidence [35], it can be inferred that determinants of OH food consumption within specific populations, like students, need to be further considered in order to inform future targeted interventions to reduce impact of unhealthy foods and drinks on public health.

Strategies for better nutrition need to target students whose lifestyle is constrained by the abundance of high energy density foods and a lack of student university restaurants or canteens, where healthier foods could be served. Well-informed choices pertaining to OH foods could help reduce energy over-consumption and improve diet quality. In parallel nutrition research needs to focus on local evidence for effective nutrition interventions and policies in order to reach a sufficient understanding among Albanian specialists and professionals. It is clear that policy interventions are required to improve the quality of diet and nutrition among university students. This research also shows that eating OH, in a broad sense, is positively associated with the likelihood of consuming less fruits and vegetable products and higher average food prices. Although difference in nutritional quality was minor, sufficient elements warrant careful monitoring of nutritional quality of overall diet.

### Study limitations and future outlooks

Although a one day 24-hours recall is appropriate to estimate population mean intakes in a crosssectional study with a relatively moderate sample size, future research should include multiple days, larger sample size and detailed information on socio-economics and demographics, as well as expanded in multiple settings and cities. However due to logistic and infrastructural constrains it was not possible to include more recalls, subjects and variables. This study was also constrained by contextual technicalities, specifically regarding the fact that there are no national food composition tables in Albania and there is very little to no research done in this field. For this reason, Greek Food Composition Tables were used, due to similarities in dieting and nutritional habits. Another difficulty, to be considered for future research is the lack of weights and measurement charts for foods and food containers in Albania. To overcome this challenge in the present study, weights and measurement charts were produced by the research team in Albania, exclusively for this study, based on which a food picture booklet was made.

Additional information on other variables like income, degree of processing of foods, residence (urban-rural), etc. would enable establishing a clearer framework for the drivers of eating OH among students in Albania, as well as a more detailed picture of the nutritional quality.

Future research may also consider conducting a national food intake survey to benchmark and increase understanding of Albanian dietary practices. Research in this regard should look at differences in nutrient content and degree of processing between AH and OH foods in general and between types of foods and drinks in order to explain changes in consumers’ nutritional status.

## Conclusion

Eating OH is common amongst the student population in Albania. Findings of this study justify the need to monitor the nutritional quality of diets, both OH and AH foods and drinks consumption and their implication for student communities’ health and beyond. Overall, there is a strong warrant for further investigation of nutritional studies in order to improve and expand knowledge on determinants of poor dietary choices and food intake.

This study provides a first assessment of current dietary habits of the studied population and can be used as baseline data for designing and conducting future studies. More detailed evidence on associations of OH and AH eating and anthropometric outcomes is needed to clearly recognize potential health implications and drivers of malnutrition in the country-specific context. We also showed the need for a coherent body of evidence and tools that are necessary for future nutritional studies of the same nature in Albania, with the final goal of supporting effective, evidence-based interventions that may reduce the impact of malnutrition on public health. A great opportunity exists within the current atmosphere of the UN Decade of Action on Nutrition (2016-2025) to address and advance nutrition narrative and end malnutrition in all its forms in Albania and in the Balkans region.

## Acknowledgements

We would like to thank our colleagues from the Department of Public Health at University of Medicine Tirana who greatly assisted the research and particularly Ms Teuta Muhollari for her valuable contribution during data entry.

## Supporting information

‘S1 Supplementary material’ can be accessed online.

